# Quantitative monitoring of neuronal regeneration by functional assay and wireless neural recording

**DOI:** 10.1101/2023.09.21.558793

**Authors:** Min-Zong Liang, Jenq-Wei Yang, Hsin Chen, Mei-Yuan Cheng, Linyi Chen

**Author notes:** **Correspondence:** Linyi Chen, PhD. Department of Medical Science, National Tsing Hua University, Hsinchu, 300044, Taiwan. Equal contribution (Min-Zong Liang, Jenq-Wei Yang).

## Abstract

Brain injury is heterozygous in nature and no single scheme is ideal for diagnosis and prognosis. Assessing proteins in cerebrospinal fluid is limited or not applicable after surgery whereas plasma biomarkers can only report occurrence of injury. The lack of regeneration markers renders the difficulty during drug discovery. This study aims to establish a potential reporter that correlates the regeneration progress of injured brain neurons. According to our recent publication, treatment of mitochondrial uncoupler carbonyl cyanide 4-(trifluoromethoxy) phenylhydrazone (FCCP) could rescue motor function deficit of mice after traumatic brain injury. We thus measured the local field potential (LFP) at sites proximal to the injury region. The recorded neuronal activity reported stimulation of right forelimb of mice. Our experimental results indicate that the recovery of evoked LFP correlates with FCCP treatment and could potentially be used as a regeneration biomarker. These promising findings suggest future application of a wireless, non-invasive recording system as a potential companion diagnosis device during drug development.

---

Neurological dysfunctions can be caused by trauma, stroke, autoimmune disease, aging, tumor, surgery, and infection. A complex set of clinical symptoms, including motor function deficit, coordination, speech, cognition, learning and memory, are associated with one or more insults. Among these, motor function deficit caused by traumatic brain injury (TBI) could significantly affect life quality, production loss and social impairment. Nonetheless, there is currently no medicine to promote neuronal regeneration after brain injury. One obstacle during drug development is the lack of regeneration biomarkers, despite several brain injury markers, S100 calcium-binding protein B and glial fibrillary acidic protein, have been proposed. Several imaging modalities are used to obtain the necessary information for patient care and prognosis. X-ray computed tomography is used as an imaging modality for assessing acute phase of TBI whereas magnetic resonance imaging is utilized in subacute and chronic cases when post-concussive symptoms are persistent. However, not all imaging modalities employed in TBI diagnosis reveal functional modifications. In addition, these two measurements cost a lot and are not accessible at local clinic. Behavior outcome and cognitive improvement have been used as measurements for drug efficacy. However, these parameters could largely depend on rehabilitation over time instead of drug administration. In addition, treating TBI patients early is expected to provide better outcome and reduce the development of neurodegenerative diseases later in their life. Thus, establishing a system to report drug effects during the acute phase of TBI is essential to facilitate drug development for treating brain injury. In this study, we used traumatic brain injury mice as a biological model, aiming to correlate neural activity and behavior improvement during neuronal regeneration. The local field potential (LFP) evoked by tactile stimulation is recorded before and after TBI. The feasibility of using the evoked LFP to reflect the functional outcome quantitatively and robustly is investigated. We shall further advance this technology toward wireless, non-invasive recording system for monitoring neuronal regeneration.

The mitochondrial oxidative phosphorylation system transfers protons from mitochondrial matrix to intermembrane space to form a proton gradient, which drives ATP synthesis (Hatefi, 1985). Mitochondrial uncoupler carbonyl cyanide 4-(trifluoromethoxy) phenylhydrazone (FCCP) induces a proton leak across the inner mitochondrial membrane to depolarized mitochondria. Nonetheless, it was reported that optimal dosage of FCCP reduced mitochondrial dysfunction and improves cognitive outcome of rat after TBI (Benz & McLaughlin, 1983; Pandya et al., 2009) . To understand the mechanism and test the effect of FCCP on motor function, we have examined the effect of FCCP on the cleavage of a mitochondrial protein phosphoglycerate mutase 5 and the correlation to neurite re-growth of injured brain neurons. As reported recently that one dose intranasal administration of 0.1 mg/kg FCCP promoted mitochondrial biogenesis and neurite re-growth of injured cortical neurons (Liang et al., 2023). For TBI mice, we have established controlled cortical impact (CCI) protocol on left brain using C57BL/6 mice and gridwalk test was performed as a behavior assessment to determine whether TBI led to foot-fault. FCCP was intranasally administered 6 hrs after CCI, numbers of foot-faults of right forelimb were calculated (Fig. 1A-C). The number of foot-faults of right forelimb of CCI mice treated with 0.1 mg/kg FCCP or DMSO (vehicle) were increased on 1 day post injury (dpi), compared to sham mice (Fig. 1D, E), indicating that motor function of right forelimb was impaired after CCI. We observed a reduction in the average number of foot-faults of right forelimb of CCI mice treated with 0.1 mg/kg FCCP on 3 dpi, which became significantly decreased by 74% on 6 dpi compared to mice treated with DMSO (Fig. 1F, G). These results indicate that 0.1 mg/kg FCCP improves functional recovery of right forelimb foot-faults after brain injury.

**Fig. 1.**
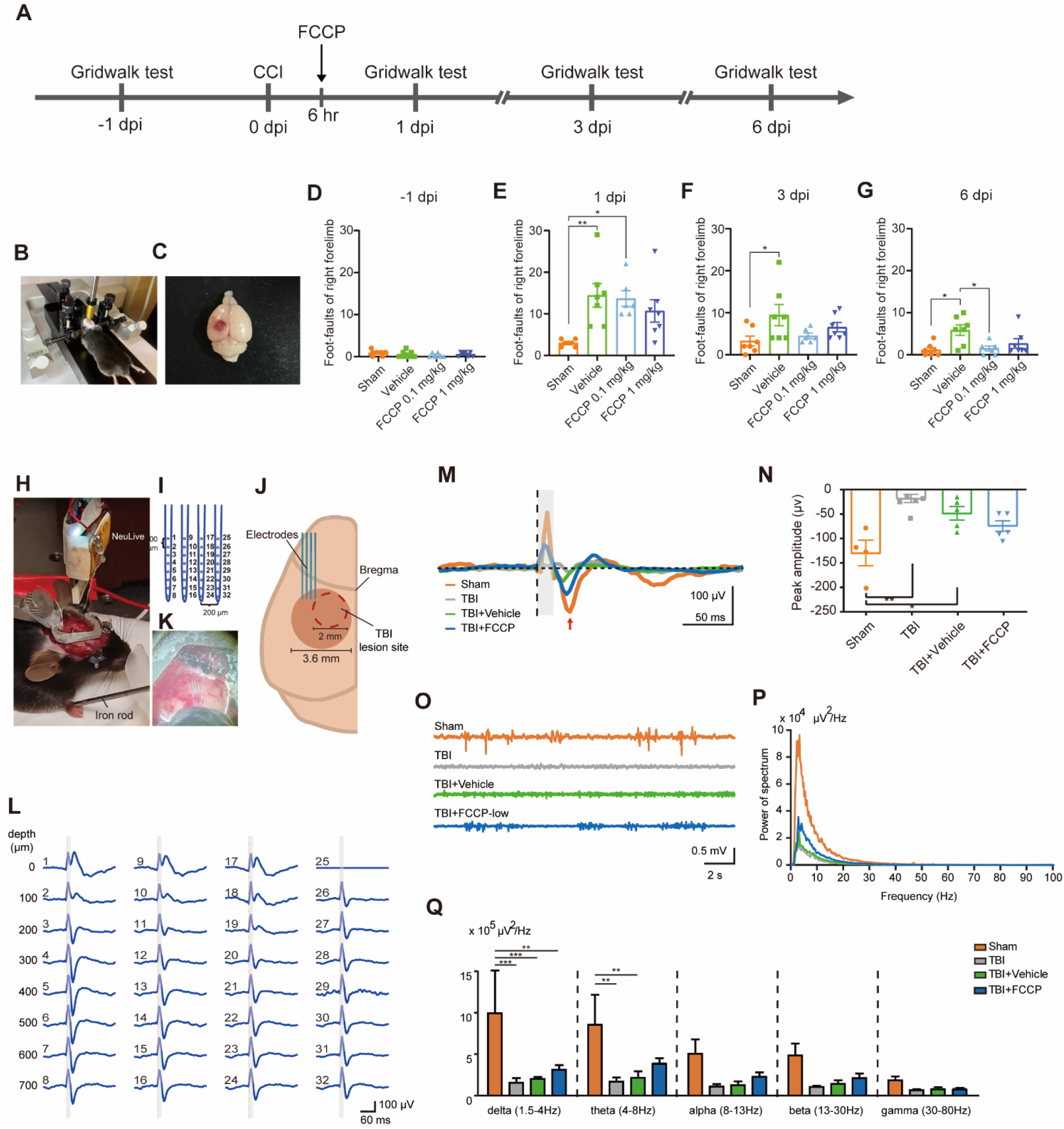
Design of multi-electrode arrays for measuring LFP and reporting brain injury. (A) Experimental diagram of the gridwalk test. FCCP or vehicle (DMSO) was nasally administrated 6 hrs after CCI. Gridwalk test was performed on -1, 1, 3 and 6 dpi to examine foot fault of CCI mice. Mice were impacted with CCI impactor. (C) Dissected brain of CCI mice. Scale bar, 5 mm. (D-G) The number of foot-faults of right forelimb of CCI mice in 5 min. Data are presented as mean ± SEM (n = 6-7 mice/group). * p < 0.05, ** p < 0.01, one‐way ANOVA with Tukey’s multiple comparisons. (H) A photo shows the experimental setup including a C57BL/6 mouse under light isoflurane anesthesia (1.2∼1.5%), a NeuLive recording head stage, a silicon based multi-electrode array for LFP recording, and an iron rod for tactile stimulation. (I) A schematic illustration of the 4-shank 32-channel multi-electrode array used in this study. (J) A top view photo shows the insertion site of the multi-electrode array. (K) A schematic illustration of the insertion site of the multi-electrode array (0.5 mm posterior and 2 to 3 mm lateral from bregma). (L) An example of the average evoked LFP by tactile stimulation on the contralateral forelimb. The gray window indicates the period of stimulation artifact. (M) The grand averages of evoked LFP responses following contralateral forelimb tactile stimulation in different experimental conditions are shown. The onset time of stimulation is indicated by a black vertical dashed line, and the gray window indicates the period of stimulation artifact. The evoked LFP response is marked by a red arrow. (N) The negative peak amplitude of evoked response is shown for different experimental groups. Data are presented as mean ± SEM (n = 4-5 mice/group). * p < 0.05, ** p < 0.01, one‐way ANOVA with Tukey’s multiple comparisons. (O) An example of 20 sec spontaneous activity for different experimental groups. (P) The averaged FFT spectra of 300 sec spontaneous activity in different experimental groups. (Q) The summation of averaged FFT power in different frequency bands. Data are presented as mean ± SEM (n = 4-5 mice/group). * p < 0.05, ** p < 0.01, *** p < 0.001, two‐way ANOVA with Tukey’s multiple comparisons.

To examine whether neuronal activity would correlate with functional recovery, we used a 4-shank 32-channel multi-electrode array to measure somatosensory evoked response induced by contralateral forepaw tactile stimulation. To this end, 4-shank/32-channel probes were into the forepaw somatosensory area in control mice (Fig. 1H-J). The location of these probes is proximal to the future injury region, which is known respond to forepaw tactile stimulation (Fig. 1K) (Auffret et al., 2018). As shown in Fig. 1L, we have demonstrated successful recording of evoked LFP following contralateral forepaw tactile stimulation. When comparing the LFP of different treatment groups, Sham, TBI only, TBI+Vehicle, and TBI+FCCP, we selected the largest evoked LFP (in the depth between 300 to 400 μm corresponding to Layer IV) for subsequent analysis (Fig. 1M). The average amplitude of sensory evoked response was significantly smaller in TBI and TBI+Vehicle groups compared to that in the Sham group, whereas the evoked response in TBI+FCCP group was closer to that in Sham control on 3dpi (Fig. 1N). The spontaneous LFP showed the same trend as the evoked LFP (Fig. 1O-P). Further analysis of frequency bands ranging from delta to gamma activity, they were reduced in TBI and TBI+Vehicle groups compared to the Sham group in all frequency. Notably, FCCP treatment showed a trend of recovery in the low frequency bands, delta and theta (Fig. 1Q). Higher frequency bands were low in all groups due to mice in anesthesia. LFP recordings (<300 Hz) majorly represent net summation of a neuronal ensemble activity surrounding the recording site related to the synchronized synaptic activity, subthreshold membrane potential and spike afterpotential (Buzsáki et al., 2012). The different frequency bands of LFP (known e.g., as delta, theta, alpha, beta and gamma) carry the information about the brain states and related to the different behavior state. The delta oscillation is associated with the deep stage 3of NREM sleep and occurs dominantly during unconscious state (Amzica & Steriade, 1998). For the animal under isoflurane anesthesia the power of low frequency oscillation increased, especially delta oscillation (Aksenov et al., 2019; Baek et al., 2022). Our data major represents the conditions when the animal is in sleep or unconscious stages.

This study used a wireless neuromodulator to record the evoked LFP, so as to monitor neuronal damage and regeneration after brain injury. During neuronal regeneration, the evoked LFP correlated with FCCP-promoted function recovery reliably. To reduce noise interferences, the recordings were performed in anesthetized mice in this study. Following this promising study, neural activities of a freely-moving TBI mice will be further recorded wirelessly with a non-invasive electrode (e.g. a flexible ECoG surface electrode array) for long-term monitoring. Such experiments would help to identify more natural, characteristic activities that indicate the rehabilitation of movement function reliably. For neuronal damage in other brain regions, the LFP could be evoked instead by electrically stimulating the upstream neurons of the injured region, and the electrical stimulation can be generated by the same neuromodulator in our study. All these studies would underpin wider applications of wireless, non-invasive neural recording and its potential utility in human study.

